# Compulsivity and impulsivity are linked to distinct aberrant developmental trajectories of fronto-striatal myelination

**DOI:** 10.1101/328146

**Authors:** Gabriel Ziegler, Tobias U. Hauser, Michael Moutoussis, Edward T. Bullmore, Ian M. Goodyer, Peter Fonagy, Peter B. Jones, NSPN Consortium, Ulman Lindenberger, Raymond J. Dolan

## Abstract

The transition from adolescence into adulthood is a period where rapid brain development coincides with an enhanced incidence of psychiatric disorder. The precise developmental brain changes that account for this emergent psychiatric symptomatology remain obscure. Capitalising on a unique longitudinal dataset, that includes *in-vivo* myelin-sensitive magnetization transfer (MT) MRI, we show this transition period is characterised by brain-wide growth in MT, within both gray matter and adjacent juxta-cortical white matter. We show that an expression of common developmental psychiatric risk symptomatology in this otherwise healthy population, specifically compulsivity and impulsivity, is tied to regionally specific aberrant unfolding of these MT trajectories. This is most marked in frontal midline structures for compulsivity, and in lateral frontal areas for impulsivity. The findings highlight a brain developmental linkage for emergent psychiatric risk features, evident in regionally specific perturbations in the expansion of MT-related myelination.

## Introduction

Structural brain development extends into adulthood, particularly so in regions that mediate higher cognition such as prefrontal cortex^1^. A canonical view is that this maturation is characterised by regional shrinkage in gray matter coupled to an expansion of white matter^2^. However, the underlying microstructural processes remain obscure. Two candidate mechanisms have been proposed^3^ namely synaptic loss (pruning) to reduce supernumerary connections, and an increase in myelination to enhance communication efficiency. Both accounts receive some support from cross-sectional and ex-vivo studies^4–7^. There are substantial inter-individual differences in these growth trajectories^8^, and the most marked changes occur within an age window where the emergence of psychiatric disorder becomes increasingly common^9,10^. This raises a possibility that psychiatric risk is tied to altered maturational brain trajectories during this critical developmental period^11,12^.

Compulsivity and impulsivity are two fundamental psychiatric dimensions^13^ and show a substantial variation in expression within a ‘healthy’ population (Supplementary Fig. 1a-e). At their extreme these features manifest as obsessive-compulsive disorder (OCD) and attention-deficit/hyperactivity disorder (ADHD) respectively. Macrostructural and crosssectional studies suggest a link to deficits in fronto-striatal regions^14–17^, but leave unanswered the question of whether compulsivity and impulsivity reflect consequences of aberrant developmental microstructural processes.

Here, we used semi-quantitative structural MRI^18^ to investigate how microstructural brain development unfolds during a transition into adulthood, and how individual variability in these developmental trajectories is linked to compulsive and impulsive traits (Supplementary Fig. 1f-g). We used a novel magnetic transfer saturation (MT) imaging protocol to provide an *in-vivo* marker for macromolecules, in particular myelin^19,20^. Importantly, MT saturation has been shown to be a more direct reflection of myelin compared to other imaging protocols, such as magnetization transfer ratio^21,22^. It also is sensitive to developmental effects^7^ which renders it ideal for tracking patterns of brain maturation within longitudinal studies involving repeated scanning of participants, a crucial feature for characterising development^23^. Using such a protocol, we show that during late adolescence and early adulthood cingulate cortex expresses the strongest myelin-related growth, both within gray and adjacent white matter. Individual differences in compulsivity are reflected in the rate of this growth in cingulate and superior frontal regions. This contrasts with impulsivity, which was associated with reduced myelin-related growth in lateral prefrontal cortex. Our results suggest that compulsivity and impulsivity traits within the healthy population may reflect a regionally specific consequence of differential unfolding of myelin growth trajectories.

## Results

### Ongoing myelin-related growth at the edge of adulthood

To assess developmental trajectories of myelin-sensitive MT, we carried out repeat scanning in 299 adolescents and young adults aged 14–24 years, up to three times in all, with an average follow-up time of 1.3±0.32 years (mean±SD) in an accelerated longitudinal design (1 scan: N=103, 2 scans: N=172, 3 scans: N=24. The sample was gender balanced and consisted otherwise healthy subjects (excluding self-reported illness a priori to avoid illness-related confounds, such as medication effects) selected to be approximately representative of the population (cf online methods for details).

Examining whole-brain maturation in gray matter revealed a brain-wide increase in myelin-related MT, with a focus within cingulate, prefrontal and temporo-parietal areas (Fig. 1a, p<.05 false-discovery rate [FDR] peak corrected; merging cross-sectional and longitudinal effects, separate effects shown in Supplementary Fig. 2a-b; mean±SD: 0.55±0.19% per year; max z-value voxel in posterior cingulate: 0.98% per year; Supplementary Table 1). This change was accompanied by increased MT in adjacent (juxta-cortical) superficial white matter, most pronounced in the exact same areas (Fig. 1b, mean±SD: 0.45±0.15% per year; max z-value voxel in posterior cingulate with 0.95% per year), consistent with the idea that connections within gray and white matter are myelinated in concert. Similar, albeit less pronounced, microstructural maturation was observed in subcortical areas such as posterior striatum, pallidum and dorsal thalamus (Fig. 1c). These findings highlight that myelin-related development in both cortical and subcortical areas is a marked feature of a transition from adolescence into adulthood, and is likely to involve both local and inter-regional fibre projections.

**Figure 1.**
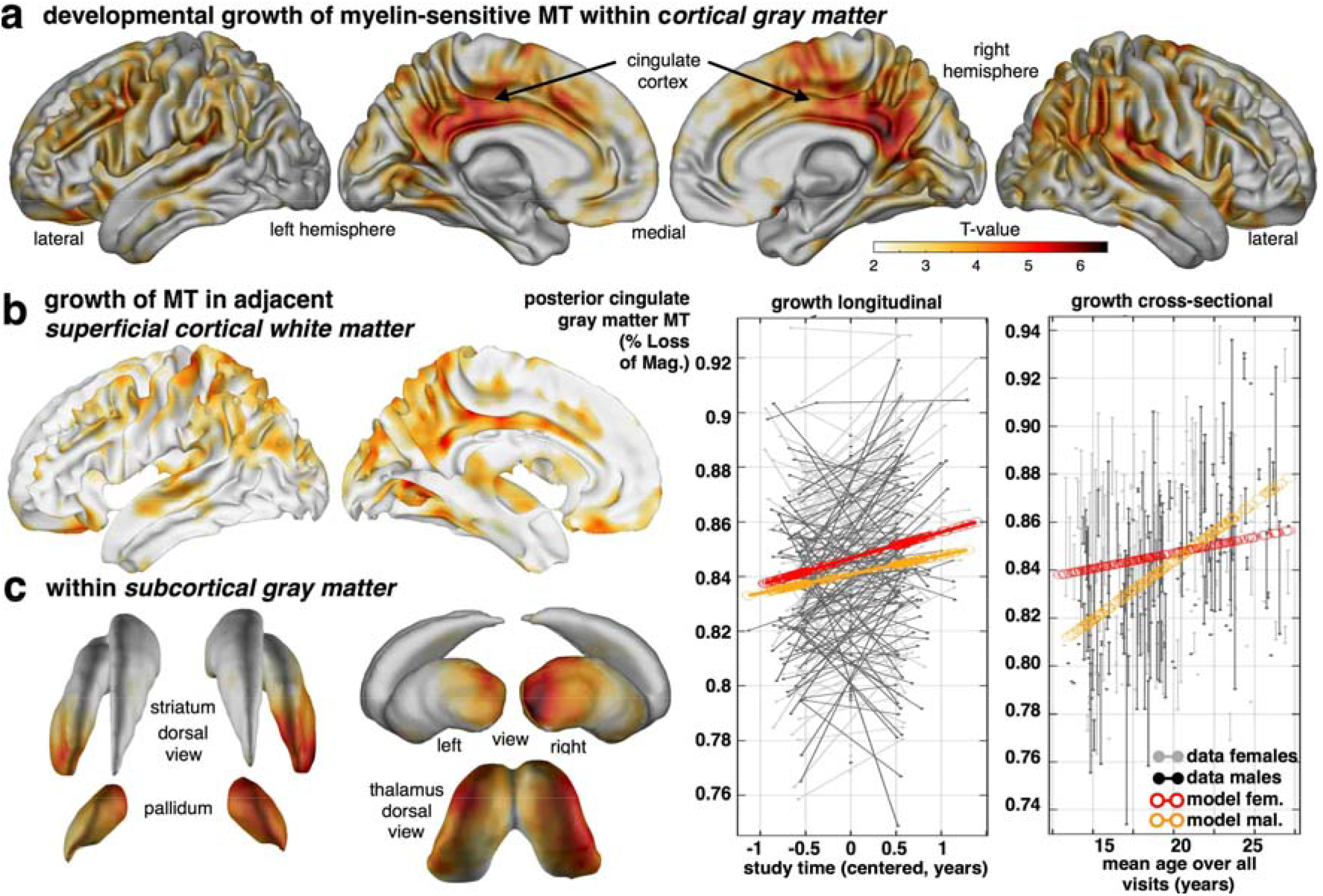
Developmental growth of myelin-sensitive MT into early adulthood. Transitioning into adulthood is characterised by profound increases in a myelin marker within cortical gray (**a**), white (**b**) and subcortical gray matter (**c**). Statistical maps of MT saturation show growth with time/visit (longitudinal) or age (cross-sectional; for specific effects of covariates, e.g. time/visit, age, sex, interactions etc., see supplementary information). (**a**) Gray matter MT growth (top row; statistical z-maps, p<.05 FDR corrected) is strongest in posterior and middle cingulate, but also present in lateral temporal, prefrontal and parietal cortex. Longitudinal model in posterior cingulate peak voxel (coloured lines in left data plot; x-axis: relative time of scan) and data (uncoloured) show an MT growth in both sexes, with a greater MT in females indicating a maturational advantage (see Supplementary Fig. 2c for region-specific sex differences, Supplementary Fig. 2d for sexual dimorphism of age-trajectories). Corresponding cross-sectional model predictions in the same region show a similar increase with age (right data plot; x-axis: mean age over visits). (**b**) MT growth in adjacent cortical white matter is most pronounced in cingulate and parieto-temporal cortex with topographical correspondence to the gray matter MT effects. (**c**) Subcortical structures express MT growth in striatum, pallidum, thalamus and hippocampus (not shown). This growth is most pronounced in posterior striatum suggesting ongoing myelin-related growth in both cortical and subcortical brain structures.

### Association between macro- and microstructural development

The observed developmental expansion of myelin-sensitive MT expressed overlapping topographies with macrostructural gray matter shrinkage (with the exception of hippocampus) and white matter expansion (Fig. 2a; Supplementary Fig. 3a-d). This raises a question as to how precisely macrostructural volume change relates to development of our myelin marker MT. A positive association in white matter volume (Fig. 2b-c; mean±SD: r=0.09±0.05) supports the notion that myelination is contributing to the observed macrostructural volume changes, as predicted by an assumption that increased myelination leads to a white matter volume expansion^24^.

**Figure 2.**
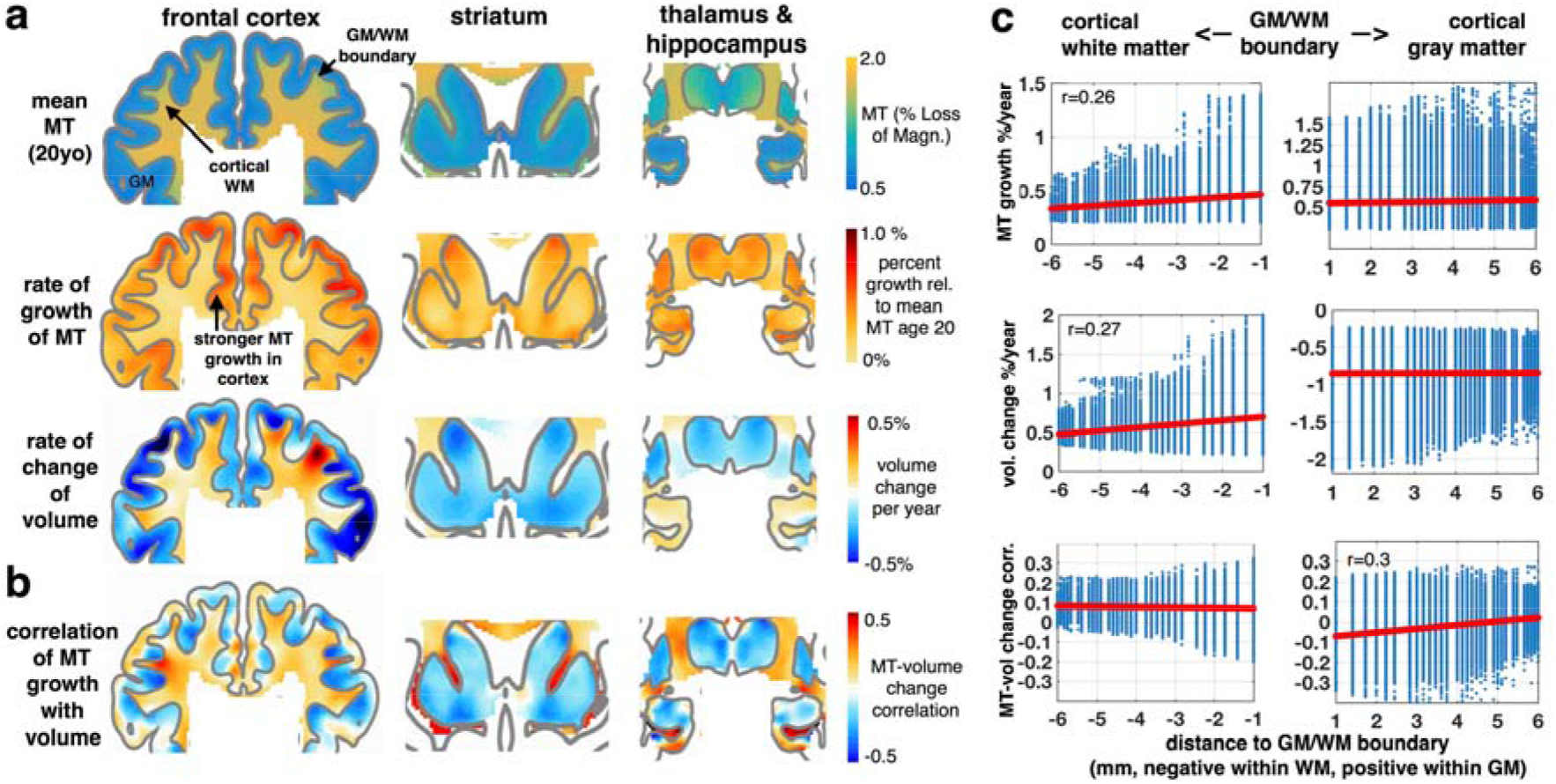
The relation between macrostructural and microstructural brain development. (**a**) Coronal sections through prefrontal (left panels), striatal (middle) and thalamus (right; MNI: y=15, 12, −14) show more myelin-related MT in white than in gray matter with a clearly preserved white-gray matter boundary (top row). Developmental change in MT (second row) shows an increase in myelin marker in both tissues, with a stronger growth in gray matter areas. Developmental change in macrostructural brain volume (third row) shows a characteristic cortical shrinkage (blue colours) in gray, but an expansion in white matter (red colours; cf Supplementary Fig. 3). Only hippocampal gray matter shows an opposite effect with continuing gray matter growth on the verge of adulthood. (**b**) Association between microstructural myelin growth and macrostructural volume change. A positive association throughout white matter supports the notion that myelination contributes to white matter expansion. In gray matter, a predominantly negative association in deep layers advocates for partial volume effects at the tissue boundary and positive associations in superficial layers (correlation was obtained from posterior covariance of beta parameters in sandwich estimator model simultaneously including longitudinal observations of both imaging modalities). (**c**) Association as a function of Euclidean distance to GM/WM boundary. Microstructural growth (top row) shows consistent myelin-related growth in both tissues, but opposite macrostructural volume change (middle row). Association between micro- and macrostructural growth is positive in white matter, independent of distance. In gray matter, the mean association changes from negative in deep layers (i.e. myelin MT change associated with reduced gray matter volume) to more positive associations in superficial layers (i.e. MT associated with a tendency to more gray matter volume).

Voxel-wise analysis in gray matter revealed a more complex association between macrostructural development and myelination (Fig. 2b-c). We observed that the association is dependent relative to where a voxel is located in the tissue. Overall consistently negative correlations (albeit relatively small) in gray matter areas close to the white matter boundary suggest that developmental myelination leads to a ‘whitening’ of gray matter, which in turn drives partial volume effects evident in a shrinkage of gray matter volume^24^. This means that gray matter volume decline in deep layers during adolescence is to some extent driven by an increase in myelination within these same areas. This negative association was found to be reduced with increased distance from the white matter boundary (Fig. 2c, bottom right panel). This suggests that ongoing myelination in superficial layers (i.e. close to the outer surface of the brain) contributes to an attenuated volume reduction and would imply that developmental macrostructural change is the result of complex microstructural processes.

### Compulsivity linked to reduced development in cingulate and striatal MT

We next asked whether individual differences in the expression of symptoms, indicative of obsessive-compulsive traits, are associated with distinct developmental trajectories in myelin-sensitive MT growth. We employed a dimensional approach and constructed a compound-score from two established obsessive-compulsive symptom questionnaires^25,26^, using the first principal component across all items (cf. supplementary information, Supplementary Fig. 1a-e). Top loading items on this score (subsequently called ‘compulsivity’) reflect compulsive behaviours, such as checking, and are tightly aligned with scores on our obsessive-compulsive questionnaires (Pearson correlations r>.8).

We focused on prefrontal cortex and striatum^14^ to examine how compulsivity relates to individual myelination over time. We found our compulsive measure was strongly linked to altered MT growth in superior lateral and medial frontal cortices (Fig. 3a, Supplementary Table 2), both in cortical gray and adjacent superficial white matter. Importantly, more compulsive subjects showed reduced MT growth compared to less compulsive subjects. A similar pattern was seen in ventral striatum and adjacent white matter (Fig. 3b). Intriguingly, the specific locations of reduced MT development were spatially circumscribed in cingulate and ventral striatum, and this regional focus closely aligns with a specific fronto-striatal loop described in primate anatomical tracing^27^ studies. This close alignment with a known anatomical circuit suggests compulsivity may be related to a deficient myelin-related developmental growth in a specific cingulate-striatal loop.

**Figure 3.**
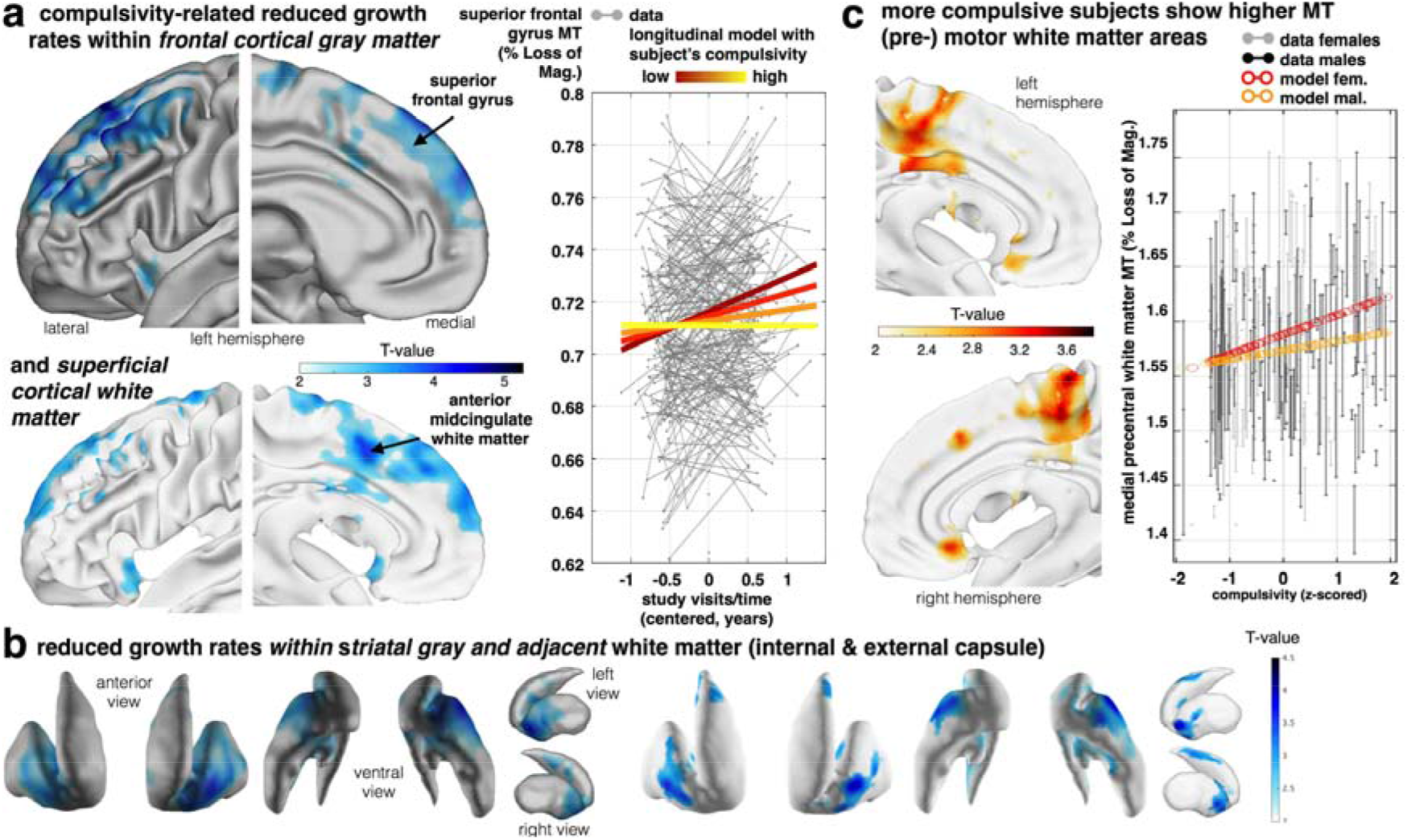
Compulsivity is related to altered fronto-striatal MT growth. Longitudinal developmental change of our myelin marker is reduced in high compulsive subjects. (**a**) Aggregate compulsivity score is related to decreased MT growth in superior frontal gyrus gray matter (upper panel; z-value=5.3, p<.0022, FDR corrected) and adjacent white matter including cingulate cortex (lower panel; blue colours depicting negative time by compulsivity interactions). Subjects with higher compulsivity scores (light yellow) compared to low scoring subjects (dark red) express significantly less MT growth over visits (coloured lines in right panel indicate the interaction effect; x-axis: time of scan in years relative to each subject’s mean age over visits). (**b**) The above slowing in cortical myelin-related growth is mirrored by a decreased developmental growth in subcortical ventral striatum (left panel) and the adjacent white matter (right panel). These findings indicate young people with high compulsive traits express slower maturational myelin-related change in a fronto-striatal network comprising cingulate cortex and ventral striatum. (**c**) More compulsive subjects showed locally increased baseline myelin marker (peak supplemental motor area, z-value=3.81, cluster FDR p<0.04,) in the medial wall. Right panel shows the plot of MT in this peak voxel over compulsivity (x-axis, z-scored) and with adjusted data (gray/black) and model predictions (red/orange, effects of interest: intercept, compulsivity, sex by compulsivity).

OCD is known to follow distinct developmental trajectories with early onset arising around the onset of puberty whereas late onset emerges at the edge of adulthood. The observed slowing of ongoing growth during adolescence is indicative of later, ongoing processes that continue into adulthood. However, this raises the question as to whether there were pre-existing myelin-related changes in our sample that would indicate aberrant myelin development prior to adolescence onset. Consequently, we investigated compulsivity main effects, and found increased myelin-sensitive MT in pre-motor, mid-cingulate, and ventromedial white matter (Fig. 3c). This is consistent with the idea that compulsive subjects have a pre-existing ‘hyper-myelination’ in these areas, similar to what is reported in rodents models of early life stress^28,29^. Importantly, these hyper-myelinated areas were generally distinct to the ones that expressed a reduced ongoing growth. This suggests that compulsivity might be linked to distinct developmental trajectories, with a pre-adolescent hyper-myelination in motor-related areas and a decreased myelination during adolescence in cingulate and frontopolar regions.

### Maturation trajectories in impulsivity

We next examined whether an impulsivity trait as assessed using questionnaire scores (Barratt impulsiveness total score) is linked to individual growth of the myelin marker in fronto-striatal areas. Importantly, in our sample we found that compulsivity and impulsivity traits were almost entirely independent (only sharing 1.4% common variance, Supplementary Fig. 1d), thus allowing us to describe their separate associations with brain development. In examining this linkage we opted to use a questionnaire measure over task-based measures of impulsivity. Our motivation for this approach arises out of the fact that the former was revealed to be more reliable, reflecting a more stable trait that is more likely to be linked to structural development. To obtain as pure a measure as possible we additionally controlled for alcohol consumption, to prevent this factor biasing impulsivity analyses^30,31^. We found that impulsivity was associated with a widespread reduction in adolescent MT growth in lateral and medial prefrontal areas (Fig. 4a, Supplementary Table 3), effects centred on inferior frontal gyrus (IFG), the precentral sulcus and insula, both within gray and white matter. We followed up on this by next assessing whether reduced myelin-related growth was also present in the striatum. We found significant associations between impulsivity and growth in MT in the dorsal striatum, more dorsal to the changes we observed for compulsivity (Fig. 4b, centred in white matter extending into gray). This suggests that while impulsivity and compulsivity are both linked to reduced myelin-related growth in prefrontal and striatal areas, these alterations occur in distinct anatomical regions (cingulate vs IFG; ventral vs dorsal striatum). Interestingly, both compulsivity and impulsivity showed a reduced growth in the anterior insula, possibly expressing a common, transdiagnostic vulnerability.

**Figure 4.**
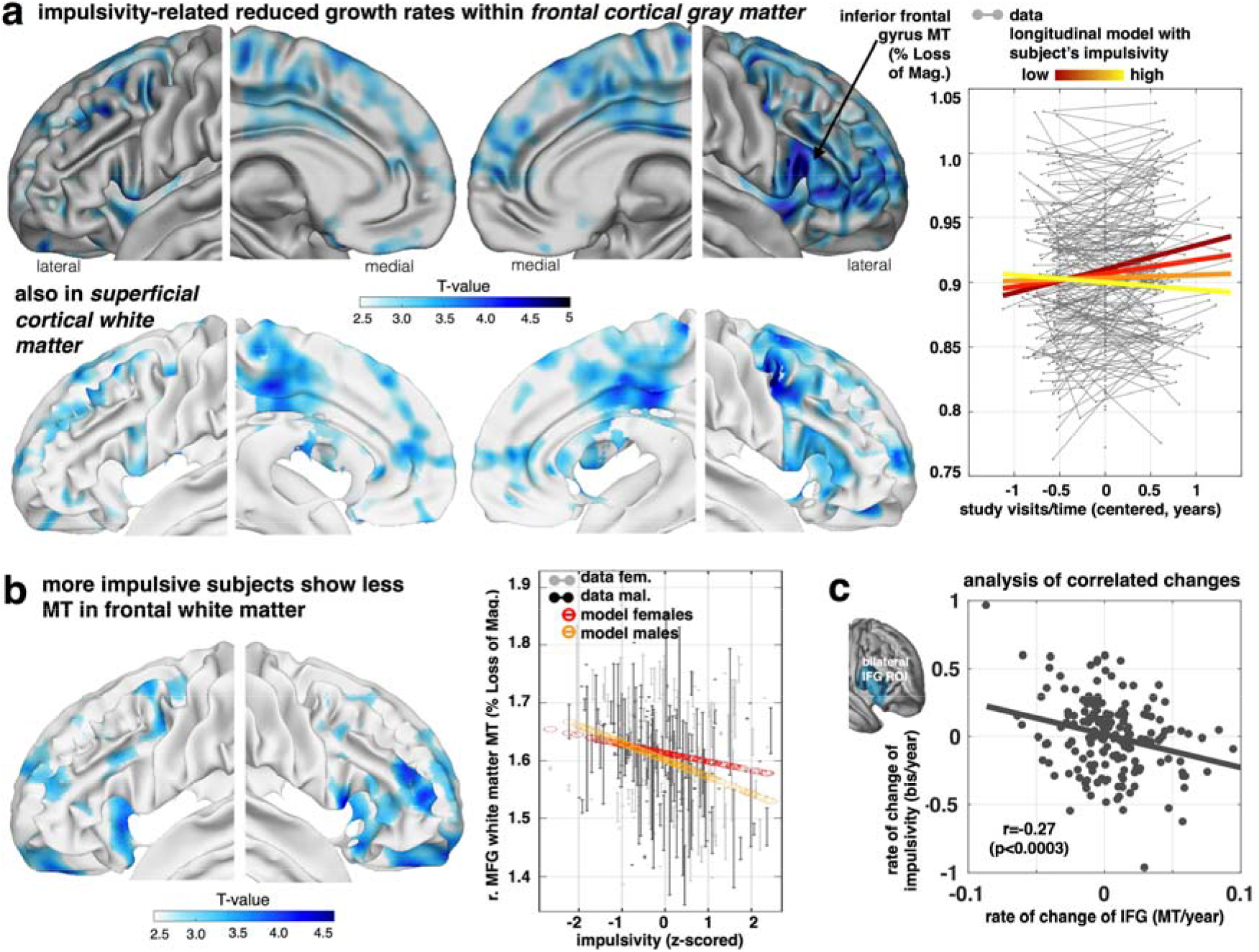
Decreased frontal growth in myelin-sensitive MT in impulsivity. Our myelin marker in frontal lobe is tied to impulsivity traits. (a) Impulsivity is associated with a reduced growth of MT in lateral (inferior and middle frontal gyrus), medial prefrontal areas (and motor/premotor and frontal pole) and anterior insula in both gray (top panel) and adjacent white matter (bottom panel) depicting negative time by impulsivity interactions (z maps, p<.05 FDR corrected). Plot shows that subjects with higher impulsivity (light yellow) compared to low scoring subjects (dark red) express significantly less MT growth over visits (coloured lines in right panel indicate the interaction effect; x-axis: time of scan in years relative to each subject’s mean age over visits). (b) More impulsive subjects showed locally decreased baseline myelin marker (peak middle frontal gyrus, z-value=4.33, p<0.027, FDR) in lateral frontal areas (fixed for other covariates, e.g. time/visits, mean age of subject, sex). Right panel shows the plot of MT in this peak voxel over impulsivity (x-axis, z-scored) and with adjusted data (gray/black) and model predictions (red/orange, effects of interest: intercept, impulsivity, sex by impulsivity). (c) Bilateral IFG not only shows a reduced myelination process for higher impulsivity (as shown in a, b), but this reduced growth rate is more strongly expressed in subjects who manifest an accentuated impulsivity growth over study visits, such that subjects who manifest an even more restricted growth in myelin become more impulsive.

As for compulsivity, we next investigated development-independent levels of myelination in impulsivity. This is important because a pre-existing ‘hyper-myelination’ would suggest a normalisation during adolescence, whereas a ‘hypo-myelination’ prior to adolescence onset would imply that a deficient myelination was further accentuated during adolescence. In contrast to compulsivity, we found a main effect of impulsivity as marked by hypo-myelination across several brain areas including IFG (Fig. 4c). Importantly, these baseline effects were found in the same areas that also showed a reduced ongoing growth. This means that for impulsivity a gap in myelination exists prior to adolescence, with this gap widening further during a transition into adulthood. The same effects were found when analysing across the entire prefrontal cortex, where a reduced MT growth reflected a risk for both psychiatric traits, but where a developmental hypo-myelination in impulsivity is further accentuated during late adolescent development (Supplementary Figure 4a-b).

Lastly, we examined whether and how MT change related to the development of impulsivity traits. Although we did not see age-related change in impulsivity across the entire group, nevertheless there was substantial variability within individuals (cf. supplementary information). We thus investigated whether myelin growth in IFG, a key region previously implicated in impulsivity^16^, was related to ongoing changes in impulsivity. We found that a change in IFG MT was negatively associated with impulsivity change (r=-.27, Fig. 4c), indicating that individuals with the least ongoing myelin growth had a worsening impulsivity over the course of the study (irrespective of other covariates, such as baseline impulsivity or age). Similar effects were also seen when conducting a voxel-wise analysis (cf. Supplementary Figure S4c).

## Discussion

Myelin enables fast and reliable communication within, and between, neuronal populations^32,33^. Using a longitudinal, repeated-measures MRI scanning design in a unique developmental sample, we provide *in-vivo* evidence that myelination continues into adulthood as evident in a pronounced myelin-related whole-brain MT growth. We find that the macrostructural growth pattern closely resembles that expressed in our myelin marker. The positive association between these measures in white matter suggests that macrostructural volume change is, at least in part, driven by myelination. In gray matter, depth-dependent associations suggest that macrostructural volume reduction in adolescence is the result of multiple microstructural processes. In superficial layers, ongoing myelination seems to attenuate the impact of a pruning effect, leading to slowing in gray matter volume decline. In deeper layers, close to the gray-white matter boundary, ongoing myelination appears to contribute to an inflated estimate of volume reduction, where a myelin-induced ‘whitening’ of gray matter can result in a misclassification of gray matter voxels (i.e. partial volume effects^24^), leading to an apparent volume reduction. This implies that developmental neuroimaging that avail of markers sensitive to specific microstructural processes can provide more accurate measures of the mechanisms underlying ongoing brain development.

Critically, we found that individual differences in myelin-related MT growth during development is linked to variation in the expression of individual psychiatric risk factors, even within this otherwise healthy sample. Both compulsivity and impulsivity were associated with a reduction in MT growth, though each is characterized by distinct spatial growth trajectories. An MT growth deficiency in compulsivity is expressed in cingulate and ventral striatum, and this contrasts with a dominant dorsal striatum and lateral prefrontal focus for impulsivity. The latter region in particular is widely implicated in impulsivity-related disorders, such as ADHD^16^, and is suggested to play a critical role in attention and response inhibition^34,35^.

In our study we opted to use a relatively broad definition of impulsivity, which suggests that developmental processes within this area may signal a general vulnerability to impulsivity. We carefully controlled for alcohol consumption, ensuring that our findings were not biased by its known effects on signatures of brain structure^36,37^. Our findings indicate that individual differences in myelin-related development are well captured by relatively broad markers for psychiatric vulnerability, though more refined cognitive endophenotypes may yield spatially more defined developmental deficits^30,31,38^. By contrast, for compulsivity, we found a specific fronto-striatal network showed decreased MT growth within cingulate and ventral striatum, areas that are prime targets for invasive treatments for OCD^39,40^. Additional ‘hyper-myelinated’ areas might form a separate, pre-adolescent risk for compulsivity (Fig. 3c). Our findings indicate that compulsivity is linked to multiple distinct aberrant myelination trajectories in anatomically discrete networks^27^.

Embracing a longitudinal developmental approach, such as the one used in this study, poses distinct developmental questions. In relation to impulsivity and compulsivity, we can ask how a stable psychiatric trait is related to longitudinal change as well as baseline myelination differences, where the latter are more indicative of influences emerging prior to recruitment into our study. In the case of impulsivity, we found that ongoing growth took place in the same regions that also showed a difference in baseline myelination, advocating for a pre-existing myelination gap that further expands during adolescence. In compulsivity, by contrast, baseline hyper-myelination affected areas largely segregate from those showing reduced ongoing growth, advocating for distinct developmental trajectories in separate brain networks. These results also imply that greater levels of myelination are not beneficial per se, but that instead it is the developmental trajectory that matters most. A deviation on either side from a normative developmental trajectory appears to carry an enhanced risk for the emergence of psychiatric symptoms.

An extension of the approach outlined above is to ask how ongoing change in psychiatric traits relate to ongoing brain maturation (i.e. correlated change). Strikingly, we found that IFG growth change was indicative of change in impulsivity. Subjects who showed worsening of their impulsivity were also those who showed the least myelin-related growth in IFG. Thus, during the transition into early adulthood even though impulsivity traits as a whole do not change at a population level, individual psychiatric risk trajectories show meaningful variation, and this in turn is reflected in specific patterns of brain maturation.

A key challenge for human neuroscience is to assess the cellular mechanisms that underlie macrostructural change *in-vivo*^41^. This assumes particular importance in developmental neuroscience where longitudinal, repeated-measures, approaches are critical for understanding brain development^23^. Our focus in this study on a magnetization transfer (MT) saturation protocol as a proxy for myelin content was based on evidence of its sensitivity to myelin and related macromolecules^18^, as well as the fact this measure is robust to instrumental biases^21^. There is also evidence for a strong relationship between MT and myelin as measured in histological studies^19,20,42^ and we also showed previously that MT is linked with myelin gene expression^7^. Our longitudinal findings extend the importance of MT as a myelin marker with relevance for individual differences, in so far as we show myelin-related effects are expressed in both white and gray matter, but more pronounced in the former as has been found in ex-vivo studies^4^. Taken together our findings suggest that MT is an important, albeit imperfect, indicator of myelin.

The transition into adulthood is a particularly vulnerable stage for the emergence of psychiatric illness^10^. Our findings suggest this expression is tied to ongoing microstructural brain development. The brain’s potential to dynamically adjust its myelination^43^, for example as a function of training^44^, points to the potential of interventions that target the specific impairments. Such interventions might offer a novel therapeutic domain to lessen a developmental vulnerability to psychiatric disorder.

## Acknowledgments

A Wellcome Trust Cambridge-UCL Mental Health and Neurosciences Network grant (095844/Z/11/Z) supported this work. RJD holds a Wellcome Trust Senior Investigator Award (098362/Z/12/Z). The UCL-Max Planck Centre is a joint initiative supported by UCL and the Max Planck Society. TUH is supported by a Wellcome Sir Henry Dale Fellowship (211155/Z/18/Z) and a grant from the Jacobs Foundation. MM was supported by the Biomedical Research Council. The Wellcome Trust Centre for Neuroimaging is supported by core funding from the Wellcome Trust (091593/Z/10/Z). First, we would like to thank Ric Davis and FIL IT support for making large sample analysis feasible and more efficient. Thanks goes also to Gita Prabhu for continuous support during course of the project. We would also like to thank specific experts for input in relation to applied and technical methods, particularly Robert Dahnke, Will Penny and Ged Ridgway, Martina Callaghan, Nik Weiskopf and Bogdan Draganski, John Ashburner and Christian Gaser, Guillaume Flandin, Tom Nichols, Bryan Guillaume, Jorge Bernal-Rusiel, Manuel Völkle, Charles Driver, Andreas Brandmeier, Fred Dick, Matthew Betts, Geert-Jan Will, and Rogier Kievit. Finally, GZ would like to thank Emrah Düzel for support at the DZNE.

## Author contributions

E.T.B., I.M.G., P.F., P.B.J., NSPN Consortium, M.M, and R.J.D. designed the experiment. G.Z., T.U.H. and NSPN Consortium performed the experiment and analysed the data. G.Z., T.U.H., U.L. and R.J.D. wrote the paper.

## Data Availability Statement

Data for this specific paper has been uploaded to the Cambridge Data Repository (https://doi.org/10.17863/CAM.12959) and password protected. Our participants did not give informed consent for their measures to be made publicly available, and it is possible that they could be identified from this data set. Access to the data supporting the analyses presented in this paper will be made available to researchers with a reasonable request to or the corresponding authors [G.Z., T.U.H.].

## Online Methods

### Study design

In total, 318 healthy adolescents and young adults were examined in this study. These subjects belonged to an MRI branch of a larger study focusing on psychiatric traits during development (NSPN study^45^, cf supplementary information). The subjects were recruited in an accelerated longitudinal study design with 6 age bins between 14 and 24 years with roughly equal number of subjects per bin, and equal gender and ethnicity (age mean (std) 19.45 (2.85) years). Subjects were recruited in London and Cambridgeshire, selected from a larger questionnaire cohort with over 2400 subjects, after screening out subjects with selfreported pervasive neurological, developmental or psychiatric disorders. We analysed 519 available brain scans from 299 healthy individuals that passed quality control. In particular, data from 103, 172, and 24 subjects with one, two or three visits per person were available, with mean (standard deviation) follow-up interval of 1.3 (0.32) years between first and last visit. The study was approved by the UK National Research Ethics Service and all participants (if <16y also their legal guardian) gave written informed consent.

### Assessing compulsivity and impulsivity

To examine the effects of compulsivity and impulsivity traits on myelin development, we collected self-report questionnaires along with the MRI assessments. As an index of impulsivity, we used the Barratt Impulsiveness Scale (BIS)^46^ total score, a well-established and calibrated measure to assess general impulsivity. To assess compulsivity, we built a composite score (using principal component analysis, cf supplementary information) from two established obsessive-compulsive questionnaires (Supplementary Fig. 1a-e, revised Obsessive-Compulsive Inventory^26^, revised Padua Inventory^25^). Compulsivity and impulsivity trait measures showed a very minor correlation r=0.119 in the large behavioural sample, supporting a notion of rather independent dimensions (less than 1.4% shared variance). Linear mixed-effects modelling (LME) revealed that both indices did not substantially change during the study period, which is why we used aggregate scores (LME intercepts) for most of the subsequent MRI analyses (cf supplementary information).

### MRI data acquisition and longitudinal preprocessing

Brain scans were acquired using the quantitative MPM protocol^47^ on three 3T Siemens Magnetom TIM Trio MRI systems located in Cambridge and London. Isotropic 1mm MT maps were collected to quantify local changes in gray and adjacent white matter and all image processing was performed using SPM12 (Wellcome Trust Centre for Neuroimaging, London, UK, http://www.fil.ion.ucl.ac.uk/spm), the h-MRI toolbox for SPM^48,49^ (https://github.molgen.mpg.de/VBQ-toolbox-group/hMRI-Toolbox-public), Computational Anatomy toolbox (CAT12, http://www.neuro.uni-jena.de/cat/) and custom made tools.

Magnetization transfer saturation (MT) maps provide quantitative maps for myelin and related macro-molecules, and correlate highly with myelin content in histological studies^19,20^. MT thus overcomes the limitations of previous methods to measure white matter integrity, such as diffusion tensor imaging, that are insensitive to attribute changes in diffusivity to actual microstructural changes such as myelin^52^ and outperforms earlier protocols such as magnetization transfer ratio^21^. This is important also because myelin patterns are defining for brain anatomy and are used for subdividing brain structures^50,51^.

To assess the microstructural myelin-related MT changes during development, we used a longitudinal processing pipeline that follows the following steps (Supplementary Fig S1f, details cf supplementary information). To normalise the images, we performed a symmetric diffeomorphic registration for longitudinal MRI^53^ and subsequently segmented each midpoint image into gray matter (GM), white matter (WM) and cerebrospinal fluid using the computational anatomy toolbox. MT maps from all time-points were then normalized to MNI space using geodesic shooting^54,55^, spatially smoothed preserving GM/WM tissue boundaries, and manually as well as statistically inspected (using sample covariance-based sample homogeneity measures, CAT toolbox). Lastly, we constructed masks for both gray and adjacent white matter using anatomical atlases for subsequent analysis (cf. illustrated in Fig. 2). As compulsivity and impulsivity are primarily associated with deficits in frontal and striatal networks^14^, we constrained the analyses of these psychiatric dimensions to striatal and prefrontal regions (cf supplementary information).

To relate these quantitative (VBQ) to more conventional metrics (i.e. Voxel-Based Morphometry), we normalized tissue segment maps accounting for existing differences and ongoing changes of local volumes using within- and between-subjects modulation. The obtained maps were spatially smoothed (6 mm FWHM).

### Longitudinal MT analyses

In this study, we employed a longitudinal observational design to explore myelin-related MT development in late adolescence and early adulthood. Traditional cross-sectional approaches employ between-subject measures to study age-related differences rather than within-subject changes. These can be affected by biases^56^, such as cohort differences^57,58^ or selection bias^59^, and typically require additional assumptions, such as (a) the age-related effect in the sample is an unbiased estimate of the group level average of individual within-subject effects or (b) all subjects change in the same way. Here, we follow recent analysis recommendations^60^, taking the advantage of the accelerated longitudinal design in which we study separately (in one joint model) (a) how the individual brain changes over time (from baseline to follow up(s)) and (b) how it varies with mean age of different subjects in the study, and their interaction. To do so, we used the accurate and efficient Sandwich Estimator (SwE)^60^ method for voxel-based longitudinal image analysis (http://www.nisox.org/Software/SwE; cf supplementary information). Similar to common cross-sectional GLM approaches, this so-called marginal model describes expected variability as a function of predictors in a design matrix, while additionally accounting for correlations due to repeated measurements and unexplained variations across individuals as an enriched error term (illustrated in Supplementary Fig S1g),

In our developmental analyses, we focused on the factors time/visits and mean age of the individual (over all visits) on whole-brain networks. Moreover, in order to investigate if and how compulsivity and impulsivity traits are related to brain trajectories and altered growth in fronto-striatal networks, we enriched the models by adding a main effect trait (compulsivity/impulsivity) as well as their interaction with change over time/visits. The latter metric allows to assess how MT growth is associated with compulsivity and impulsivity trait (e.g. lower MT growth in high compulsives), whereas the former indicates how a trait relates to overall MT differences across individuals, independent of all other covariates (time, mean age of a subject over all scans, sex, etc.). Unless mentioned otherwise, all analyses were performed in a dimensional way, using the subjects’ trait scores. For illustration purposes, we subsequently used group-splits to visualise models and data.

Notably, in addition to including effects time/visit, mean age of subject (further denoted age_mean), and compulsivity/impulsivity traits, all presented models were tested for indications of effects of (a) other relevant demographic factors, especially sex and socioeconomic status; (b) non-linearites (accelerations/deceleration) of brain changes (across the study age range) and of age-related trajectories, especially using time by age_mean interactions, and quadratic/cubic effects of age_mean; and (c) all first order interactions among all previous covariates (cf. supplementary information for more details). No indications for substantial non-linearities were observed for myelin-sensitive MT (cf Fig. S2e) but for volumes (cf Fig. S3c). Demographic covariates and confounds (total intracranial volume, scanner, socioeconomic status) were included in all models, and additional interactions of covariates were included when showing significant effects. This is intended to account for potential confounding effects of residual head size variations induced by tissue-weighted smoothing of quantitative MT analysis and during morphometric analysis. Additionally this allows using a consistent design (and power) across modalities. We controlled for the False Discovery Rate (FDR) during corrections for multiple comparisons in all image analyses. All correlations were reported without p-values in accordance with the recommendations from the American Statistical Association^61^, but given the sample size, the reader rests assured that these would be significant if tested.

We additionally tested for correlated changes in the supplementary material, investigating how a change in compulsivity/impulsivity relates to ongoing myelin-related changes (cf. Supplementary Information for details).

### Macrostructural changes

To be able to relate the findings from our microstructural myelin marker (MT) to traditional macrostructural markers (GM/WM volume), we performed analogue analyses (using Voxel-based Morphometry, VBM^62^) as described above on traditional normalized tissue segment maps. To quantify how developmental changes of macro-to microstructural parameters correlate, we specified a multi-modal SwE model including all volumetric and MT scans in a joint (blockdiagonal) design matrix with all covariates separately for each modality. Developmental effects within each modality are defined by respective *time/visit* and *age_mean* beta estimates of those regressors of the design matrix. After SwE model estimation, the posterior covariance of these beta parameters from volume and MT modalities were calculated and transformed into correlation (see Fig. 2b).

### Assessing wide-spread effects of compulsivity and impulsivity

To assess the effects of development and compulsivity/impulsivity on myelin-sensitive MT across the entire frontal lobe (GM, WM separately), we used linear mixed-effects modelling (LME, cf supplementary information). Besides assessing the effects of *time/visit* and *time* by (continuous) *trait* interactions, we calculated the model predictions over the study period while accounting for variations of mean age across individuals^63^. Random-effect intercepts were included and proved optimally suited using likelihood ratio tests. To assess myelin marker differences at baseline we calculated t-statistics (and p-values) of main effect of traits.

### Analysis of correlated changes of brain and impulsivity

To address whether MT development was related individual changes in impulsivity, we conducted a hypothesis-driven analysis of the bilateral IGF (anatomically defined). This LME analysis provides information about whether changes in impulsivity also reflect how quickly this brain region myelinates during the study period. The LME model used IFG MT, rates of change in IFG MT, time, their interaction, as well as the above mentioned covariates as fixed effect to predict the dependent variable impulsivity score. We visualize the observed correlated changes using simple correlations. In addition, we conducted exploratory voxel-wise correlated change analyses (cf. supplemental information).

### Code availability

Custom made SPM pipeline code for longitudinal VBM and VBQ processing is provided along with the manuscript (https://github.com/gabrielziegler/gz/tree/master/nspn_mpm_prepro_code_and_example).

